# Cysteine *S*-acetylation is a post-translational modification involved in metabolic regulation

**DOI:** 10.1101/2024.05.21.595030

**Authors:** E. Keith Keenan, Akshay Bareja, Yannie Lam, Paul A. Grimsrud, Matthew D. Hirschey

## Abstract

Cysteine is a reactive amino acid central to the catalytic activities of many enzymes. It is also a common target of post-translational modifications (PTMs), such as palmitoylation. This long-chain acyl PTM can modify cysteine residues and induce changes in protein subcellular localization. We hypothesized that cysteine could also be modified by short-chain acyl groups, such as cysteine *S*-acetylation. To test this, we developed sample preparation and non-targeted mass spectrometry protocols to analyze the mouse liver proteome for cysteine acetylation. Our findings revealed hundreds of sites of cysteine acetylation across multiple tissue types, revealing a previously uncharacterized cysteine acetylome. Cysteine acetylation shows a marked cytoplasmic subcellular localization signature, with tissue-specific acetylome patterns and specific changes upon metabolic stress. This study uncovers a novel aspect of cysteine biochemistry, highlighting short-chain modifications alongside known long-chain acyl PTMs. These findings enrich our understanding of the landscape of acyl modifications and suggest new research directions in enzyme activity regulation and cellular signaling in metabolism.

## INTRODUCTION

Protein modifications are central to cell biology across all organisms, enabling chemical diversity beyond that of the canonical amino acids and altering protein functions. These modifications, most often occur on amino acid side chains and are crucial for the dynamic regulation of protein chemistry and function. The discovery and functional analysis of PTMs, significantly propelled by mass spectrometry and computational tools, have revealed over 500 distinct modifications,^1^ highlighting their complexity and vital role in cellular processes and human health.

Short-chain acyl PTMs, particularly lysine acetylation, are well-known and regulate a myriad of cellular functions. Lysine acetylation can be catalyzed enzymatically or driven non-enzymatically.^2^ This potential for non-enzymatic modification has likewise been demonstrated for several acyl-CoA moieties beyond acetylation.^3,4^ One mechanism of non-enzymatic lysine acetylation explored *in vitro* involves non-enzymatic modification of the side chain of a proximal cysteine followed by transfer of the acetyl group to lysine by intramolecular transfer.^5^ While this has only been shown *in vitro* using synthetic peptides and membrane proteins incubated with acetyl-CoA, it hints at the possible presence of short-chain acyl modifications to cysteine *in vivo*. Furthermore, the relationship between acyl-CoAs and protein acylation suggests many PTMs await discovery based on these principles.

Given cysteine’s reactivity at physiologic conditions and the emerging paradigm that acyl-CoA species can induce non-enzymatic protein modifications, we set out to test for cysteine-linked protein modifications. To accomplish this, we developed a sample preparation protocol that would allow us to identify short-chain acyl modifications to cysteine and ultimately uncover a previously unknown cysteine acetylome.

## RESULTS

### Identification and confirmation of cysteine S-acetylation

To identify whether acetyl-cysteine (Figure 1A) was present in an *in vivo* setting, we performed label-free non-targeted mass spectrometry (MS) analysis of the mouse liver proteome from freshly harvested C57/Bl6J wild-type mouse liver samples without any chemical enrichment. The inherent lability of an S-linked short-chain acyl PTM rendered standard protocols for preparing tissue samples for MS analysis unusable for these studies because conditions would degrade a putative acetyl-cysteine modifiation. The thioester bond, which connects acyl modifications to the cysteine sidechain, is sensitive to temperature, pH, and reducing agents and is susceptible to nucleophilic attack, which degrades short-chain cysteine acyl PTMs over time.^6^ To address this, we developed a system for MS sample preparation that preserves the cysteine acetylome. Incubation with tris(2-carboxyethyl)phosphine (TCEP), which is minimally reactive towards the thioester bond, allows for the preservation of the cysteine acetylome compared to other common reducing agents such as dithiothreitol (DTT).^7^ Sample pH was kept at or below 7 using a modified version of the commercially available S-trap sample preparation system.^8^ Crucially, the S-trap system, along with extensive optimization, allowed us to streamline the sample preparation process to as little as 5 hours from the initial reconstitution of a frozen pulverized tissue sample into lysis buffer to elution of tryptic peptides from solid phase extraction (SPE) and sample drying by speed vac. Using data-dependent acquisition, peptide samples were analyzed by reversed phase nanoflow LC and tandem MS (MS/MS). Raw data was searched using cysteine acetylation and lysine acetylation as variable modifications. Initial searches revealed over 400 sites of cysteine side-chain acetylation in the mouse liver proteome. Out of 25,438 total unique peptides identified in these MS analyses, 463 contained putative cysteine side-chain acetylation. These acetyl-cysteine-containing peptides represented ∼8.4% of the 5,512 cysteine-containing peptides identified. By comparison, 384 peptides bearing lysine acetylation modifications were identified from these same samples. These results reveal the cysteine acetylome is approximately as extensive as the lysine acetylome but has been previously undetectable likely due to sample preparation methods.

**Figure 1.**
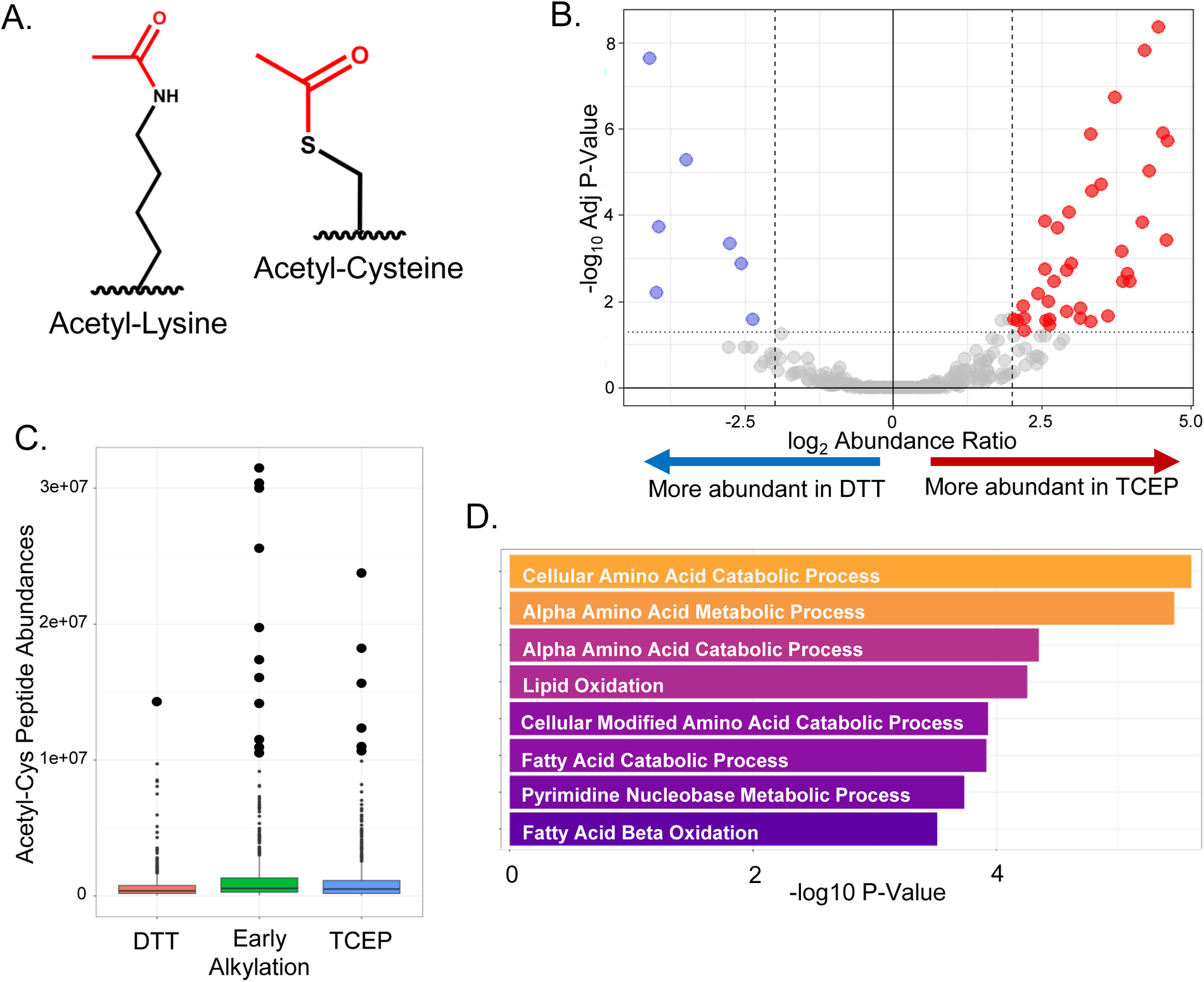
Identification of the cysteine acetylome by MS analysis and ablation of the cysteine acetylome by reduction with DTT. A. Structures of Acetyl-Lysine and Acetyl-Cysteine. B. Volcano plot showing relative abundance of sites of cysteine acetylation in mouse tissue following reduction with TCEP compared to reduction with DTT. Cutoffs at 2x fold-change and p<0.05. C. Distribution of peptide abundances of all cysteine acetylated peptides from initial preparation compared with early rapid alkylation preparation. Peptides from initial prep are separated according to reducing agent (TCEP or DTT). Abundances greater than 1e+07 enlarged for emphasis. D. Pathway overrepresentation analysis of cysteine acetylated proteins identified from mouse liver samples prepared using rapid early alkylation protocol. Overrepresentation analysis was performed against all proteins identified in the sample as a background. Top eight enriched pathways shown.

To probe the chemical lability of cysteine *S*-acetylation, we tested the reactivity of the observed PTM by differentially treating samples with reducing agents and quantitatively monitoring the result. Samples were treated with either TCEP or DTT during preparation. TCEP reduces disulfide bonds without disturbing the thioester bond of acetyl-cysteine, whereas DTT reduces disulfide bonds and thioester bonds, thereby removing acetyl-cysteine modifications. ^9,10^ The proteomic workflow was coupled with label-free quantitation to monitor reaction progress. In the presence of cysteine acetylation, DTT would be expected to decrease the signal of acetyl-cysteine observed by MS. Indeed, upon comparing results from the TCEP condition and the DTT condition, a marked reduction of the acetyl-cysteine signal in the DTT condition was observed (Figure 1B). Of the acetyl-cysteine peptides that showed a statistically significant change (p<0.05), 89% showed ablation under the DTT condition. presence of a DTT-labile thioester bond linking the acetyl-modification to the side-chain of cysteines. Furthermore, this result provided chemical confirmation of the structure matching the observed mass shift on cysteine, providing further evidence that the PTM identified was indeed cysteine S-acetylation.

S-acetylcysteine is a labile modification likely formed by the cysteine side-chain’s deprotonated thiol attacking an acetyl-CoA molecule’s thioester bond.^5^ The resulting S-acetyl cysteine itself contains a thioester bond connecting the cysteine residue to the acetyl moiety. Unmodified cysteines would thus be expected to perform nucleophilic attack on acetylated cysteines, resulting in intramolecular transfer of the acetyl group over to the formerly unmodified cysteine residue. In cases of lysis under denaturing and/or reducing conditions (such as during sample preparation for proteomic analysis) this would enable broad-scale intramolecular transfer due to the sudden exposure of many unprotected cysteine residues. To test this idea, we compared samples that were immediately alkylated upon thawing with samples that had been incubated at RT for ∼1 hour before the addition of alkylating agent. We discovered that delaying alkylation in the sample preparation conditions led to a more diffuse and stochastic cysteine acetylome (Table 1 and Table 2) consistent with broad intramolecular transfer. In contrast, rapid, immediate alkylation allowed for consistent identification and higher abundance acetyl-cysteine peptide hits and prevented many low abundance, quantitatively inconsistent acetyl-cysteine hits. Indeed, the number of cysteine acetylated peptides decreased, with roughly half as many peptides identified as cysteine acetylated with immediate alkylation. Interestingly, the total number of cysteine-containing peptides identified also decreased, likely due to other thioester-containing modifications to cysteine, including known long-chain acyl modifications such as palmitoylation, which would be expected to be better preserved under these conditions, but were not considered in search for the data. Under these acetyl-cysteine-preserving conditions, MS analysis revealed 218 peptides with cysteine side-chain acetylation out of 32,314 unique peptides identified in the mouse liver proteome. In all, 4,649 cysteine-containing peptides were identified, yielding a modification rate of ∼4.7% of cysteine-containing peptides.

While the number of unique acetyl-cysteine peptides decreased, the highest abundance of acetyl-cysteine hits were more abundant (Figure 1C) and hence more consistently identified and reliably measured. This increased consistency allowed for the identification of relatively high abundance sites of cysteine acetylation, as estimated by the number of peptide spectral matches (PSMs) identified for those sites. As shown in Table 3, top hits (as ranked by number of PSMs) in the liver cysteine acetylome included catalase, pancreatic triacylglycerol lipase, glutathione-S-transferase, and glyceraldehyde-3-phosphate dehydrogenase (GAPDH). Thus, this key sample preparation step provides a more consistent and accurate picture of the cysteine acetylome.

This mapping of the cysteine acetylome provided an overview of the distribution and frequency of cysteine acetylation across the mouse liver proteome. Thus, we performed an overrepresentation analysis comparing S-acetylated proteins against all proteins identified in our experimental dataset to determine which biological pathways were most prominent in the cysteine acetylome. This analysis revealed a strong metabolic signature in the cysteine acetylome. Cellular Amino Acid Catabolic Process and Alpha Amino Acid Metabolic Process were the most enriched pathways, each with a p-value < 0.01 and a false discovery rate (FDR) < 0.05 (Figure 1D). This metabolic signature is consistent with the cysteine acetylated protein datasets containing several key metabolic proteins, including GDH, mitochondrial trifunctional protein, and GAPDH (Table 3). These results demonstrate the presence of cysteine S-acetylation occurring in mouse liver tissue *in vivo*.

### Acetyl-CoA treatment increases cysteine acetylation

Next, we tested whether acetyl-CoA could serve as a carbon source for this new modification. Given that acetyl-CoA is the canonical source of lysine acetylation^1,4^, we reasoned that acetyl-CoA incubation with liver lysate would lead to an increase in cysteine acetylation. Mouse liver lysate samples were incubated with 1 or 10 mM Acetyl-CoA at room temperature for 1 hour. As predicted, incubation with acetyl-CoA led to increased levels of total cysteine acetylation in a concentration-dependent manner (Figure S1A). Incubation with 1 mM acetyl-CoA resulted in a 3.7-fold increase in the summed intensity (*i.e.* area under the curve from MS1 extracted ion chromatograms) of cysteine acetylated peptides normalized to total peptide signal while the number of unique acetylated peptides detected only increased by 13%. When the concentration of acetyl-CoA was increased to 10mM, the summed intensity of cysteine acetylated peptides normalized to total peptide signal increased 8.6 fold over control, and the number of unique acetylated peptides increased by 60%. From this observation, we conclude that 1 mM acetyl-CoA increased the occupancy of cysteine acetylation at sites that were already acetylated in other proteins, while 10 mM increased new sites of acetylation. Interestingly, this increase in cysteine acetylation was not evenly distributed across the cysteine acetylome (Figure S1B). While most modification sites showed increased levels of cysteine acetylation, some sites showed little to no change. For example, catalase showed a sharp increase cysteine acetylation on Cys393 and Cys460 in both the 1 mM acetyl-CoA and 10 mM acetyl-CoA conditions. Other proteins such as glutamate dehydrogenase (GDH) showed a linear response to increasing concentrations of acetyl-CoA, with a 4.5-fold increase in acetylation of Cys112 upon incubation with 1 mM acetyl-CoA and a 36-fold increase upon incubation with 10 mM acetyl-CoA. GAPDH (S4R1W1 isoform) showed no increase in active site cysteine acetylation at 1mM acetyl-CoA, but a 15-fold increase upon incubation with 10 mM acetyl-CoA (Table 5). These results imply that cysteine acetylation is not randomly distributed but rather occurs differently at different sites of modification. Enzymatic activity, as well as the solvent accessibility and microenvironment (*i.e*., pKa) of a cysteine side-chain, may play a role in regulating how specific sites are modified. At supraphysiolic concentrations of acetyl-CoA (10 mM), it seems that a less targeted mechanism of cysteine modification becomes more prominent, possibly due to widespread non-enzymatic modification of cysteine (Figure S1C). Altogether, these results support a role for acetyl-CoA as the carbon donor of cysteine acetylation.

### Subcellular Localization of the Cysteine Acetylome

Because lysine acetylation has strong sub-cellular distribution patterns,^11,12^ we measured the subcellular distribution of target proteins. The strong overrepresentation of metabolic pathways in the early cysteine acetylome suggested the possibility that the cysteine acetylome could be localized to metabolically active compartments of the cell, such as mitochondria. To test this, freshly harvested mouse liver samples were fractionated by differential centrifugation into nuclear, mitochondrial, and cytoplasmic fractions. Each fraction was then prepared for MS analysis using the protocols developed earlier. To our surprise, the cytoplasmic fraction had the highest levels of cysteine acetylation, measured by the number of sites identified or by the summed intensity of cysteine acetylated peptides. The nuclear fraction showed the lowest levels of overall cysteine acetylation (Figure 2A).

**Figure 2.**
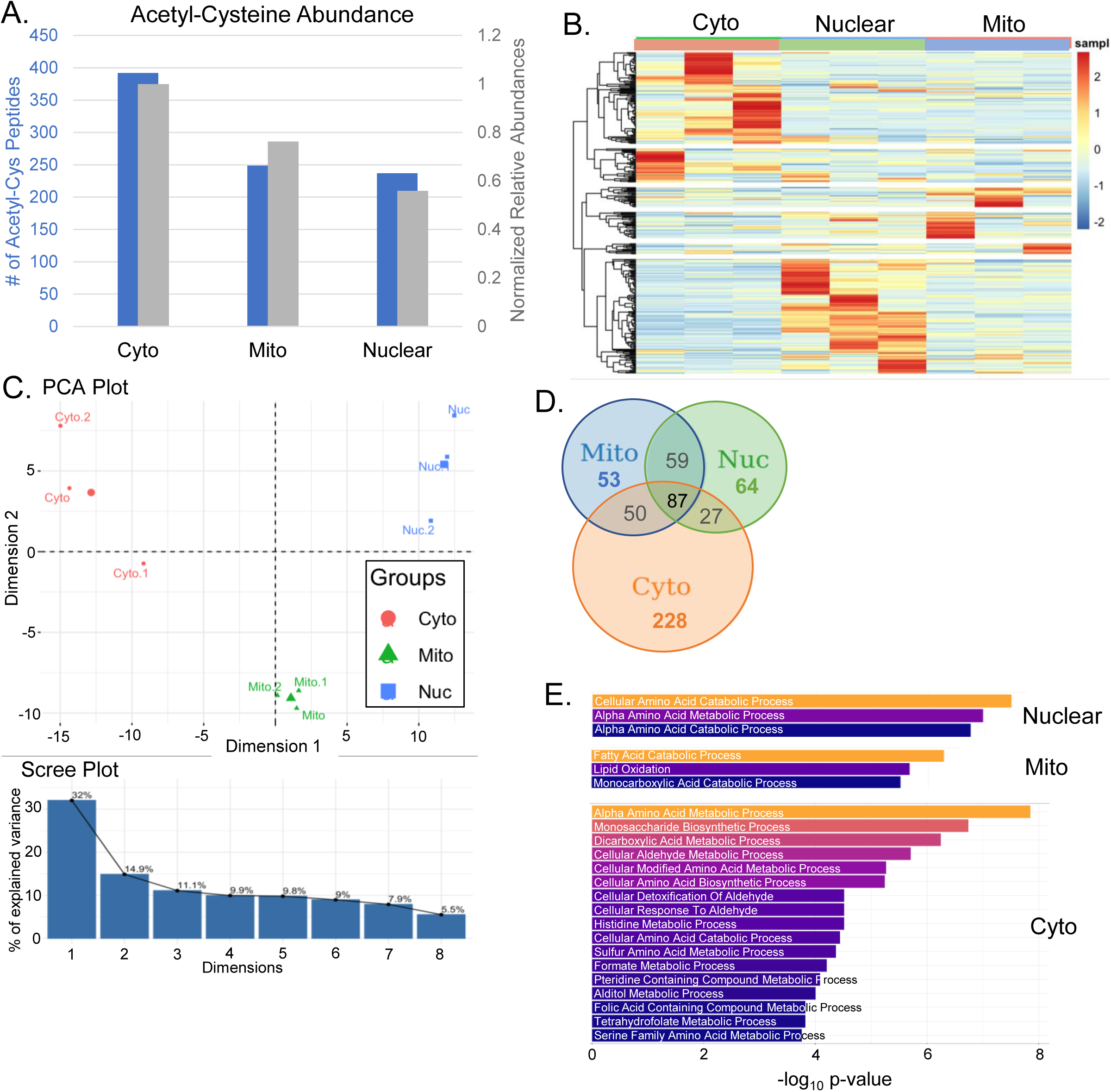
Analysis of the cysteine acetylome following subcellular fractionation of mouse liver into cytoplasmic, mitochondrial, and nuclear fractions. A. The summed intensity of cysteine acetylated peptides normalized to total peptide signal (in gray), and the number of unique S-acetylated peptides identified (in blue) for each subcellular compartment. B. Heatmap showing distribution of cysteine acetylome in each subcellular compartment. C. PCA of the cysteine acetylome in cytoplasmic, mitochondrial, and nuclear subcellular compartments. Scree plot showing the relative contribution of each of the first 10 components. D. Ven diagram showing extent of overlap of the cysteine acetylome between subcellular compartments as measured by shared sites of cysteine acetylation. E. Pathway overrepresentation analysis of the cysteine acetylome for each compartment. Showing pathways with p-value < 0.01, adj. p-value < 0.05. Pathway overrepresentation analysis was performed using cysteine acetylated genes for each compartment against the list of all genes identified in that compartment as background.

In addition to differences in overall levels of acetylation, each subcellular compartment showed distinct patterns of cysteine acetylation (Figure 2B, Figure S2). Primary component analysis (PCA) also showed separation of the cysteine acetylome for each subcellular compartment. This indicated that the pattern of the cysteine acetylation on proteins was distinct in different subcellular compartments. The first and second components of the PCA accounted for 32% and 14.9% of the observed differences, respectively (Figure 2C).

In all, 392 acetyl-cysteine-containing peptides were identified in the cytoplasmic fraction, 237 in the nuclear fraction, and 249 in the mitochondrial fraction. Only 87 peptides were shared between all three compartments (Figure 2D). The summed intensity of cysteine acetylated peptides also followed a similar trend for the three compartments (Figure 2A). Subcellular fractionation resulted in more acetyl-cysteine peptides being identified compared to unfractionated samples. This may have been due to the identification of lower abundance hits masked by the greater number of peptides injected when whole cell lysate was used. Additionally, the greater length of time prior to alkylation may have allowed for intramolecular transfer of acetylation.

Overrepresentation analysis was performed on the proteins identified in each sub-cellular fraction to identify possible pathways enriched with protein cysteine acylation. The nuclear and mitochondrial fractions each only showed three enriched pathways (p<0.01, FDR<0.05), while the cytosolic fraction showed 17 enriched pathways (Figure 2E). In all cases, metabolic pathways were overrepresented, suggesting cysteine acetylation could influence metabolic processes.

### Multi-tissue cysteine acetylome

Previous experiments had all used mouse liver tissue as a source of acetyl-cysteine-containing peptides. Thus, we performed a multi-tissue analysis to fully explore the cysteine acetylome across murine tissues. Brown adipose tissue (BAT), brain, heart, kidney, liver, lung, pancreas, skeletal muscle (Skm), and spleen samples were harvested from male, three-month-old C57/Bl6J wild-type mice. These samples were processed for MS analysis, as performed above. While the initial characterization of the cysteine acetylome in the liver focused on fully identifying all possible sites of cysteine acetylation, this multi-tissue survey focused on identifying the top sites of cysteine acetylation across a broad range of tissues. The study was performed with triplicate biological samples, and each sample was analyzed by MS once for three runs per tissue type. Surveys of the hepatic cysteine acetylome had as many as 18 runs, which allowed for deeper characterization of the cysteine acetylome but was highly resource-intensive. Because we predict higher abundance sites of cysteine acetylation are more likely to be biologically relevant, we focused on identifying these primary modification sites in this multi-tissue survey.

Each tissue showed a distinct pattern of cysteine acetylation (Figure 3B). A study QC pool was created by combining equal amounts of each tissue peptide digest, which was run in five technical replicate injections, interspersed with individual tissue sample runs. This provided an empirical average of the proteomes of all the tissues, and allowed for assessment of technical variability—yielding an average CV for all peptides measured. The study QC pool also assisted with the chromatographic alignment of features.

**Figure 3.**
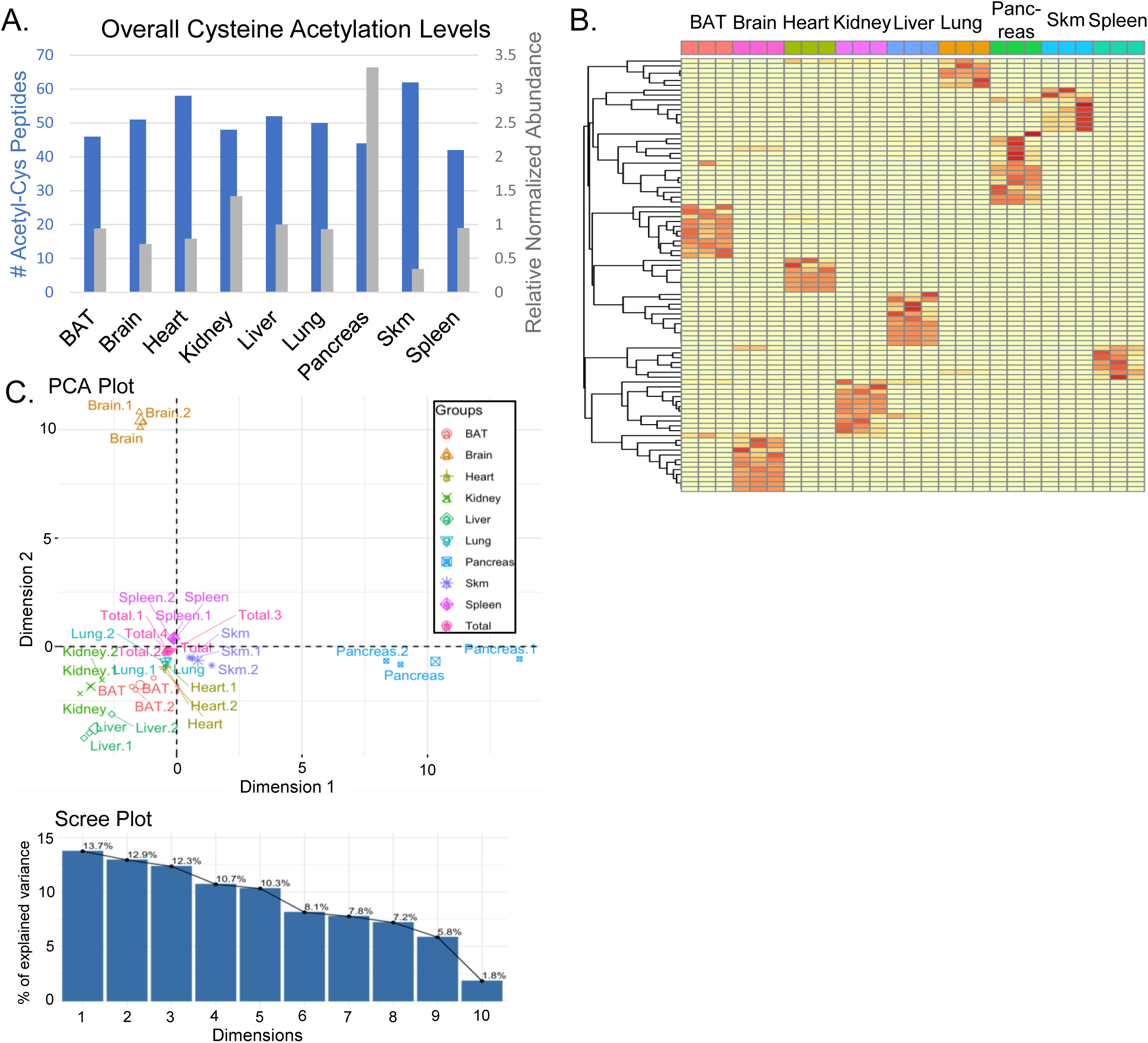
Multi-tissue survey of the murine cysteine acetylome. A. The summed intensity of cysteine acetylated peptides, normalized to total peptide signal, (in gray) and the number of unique S-acetylated peptides identified (in blue) for each tissue type. B. Heatmap showing the distribution of the cysteine acetylome for different tissues. C. PCA of the cysteine acetylome across all tissue types studied. Pooled QC sample labeled as Total. Scree plot showing the relative contribution of each of the first 10 components.

PCA showed separate acetylation patterns in each tissue type (Figure 3C). Brain, skeletal muscle, and heart showed the greatest separation, with other tissues clustering near the study QC pool labeled “Total”. The variability between individual tissue samples was slightly greater than between runs of the study QC pool sample, indicating biological variability in the cysteine acetylome beyond that inherent in the MS. Some tissues such as the heart did not have greater inter-sample variability than the study QC pool, indicating lower levels of biological variation in the cysteine acetylome for those tissues. Overall, the first and second components of the PCA accounted for 13.7% and 12.9% of the total difference observed.

Each tissue had a unique set of top cysteine acetylated proteins (Table 5). Skeletal muscle showed cysteine acetylation of Actin, and liver showed acetylation of GAPDH. Interestingly BAT, liver, spleen, and pancreas all showed cysteine acetylation of pancreatic triacylglycerol lipase. The only acetylated protein shared across all datasets was the ribosomal protein RPL4.

Overall, the levels of cysteine acetylation varied across the tissue types studied. Interestingly, the number of cysteine acetylation sites and the total normalized abundance of S-acetylated peptides did not always correlate, suggesting that the cysteine acetylome was more narrowly localized in some tissues than others. Skeletal muscle had the greatest number of cysteine acetylated peptides, while the spleen had the lowest number of cysteine acetylated peptides. The pancreas had the highest summed intensity of cysteine acetylated peptides normalized to the total peptide signal at 3.3x compared to the levels observed in the liver. Skeletal muscle had the lowest out of all the tissues tested at 0.3x that of the liver (Figure 3A).

### Effect of cold challenge on the cysteine acetylome in brown adipose tissue

To determine if cysteine acetylation changes under physiological conditions, we investigated brown adipose tissue (BAT). BAT is a highly metabolically active tissue with marked upregulation of metabolic activity when cold-challenged.^13,14^ To determine if cysteine acetylation changes during cold response, BAT was harvested from mice that were housed at RT and mice that were cold adjusted by housing at 6□C for 4 weeks. We collected BAT, processed the samples with our standard method, and characterized the cysteine acetylome by MS. We observed that levels of cysteine acetylation were largely equivalent between the two conditions (Figure 4A), but the distribution of the cysteine acetylome changed upon cold treatment (Figure 4B). PCA showed separation of the 6□C and RT samples with the first component of the PCA accounting for 65.6% of the variability observed (Figure 4C).

**Figure 4.**
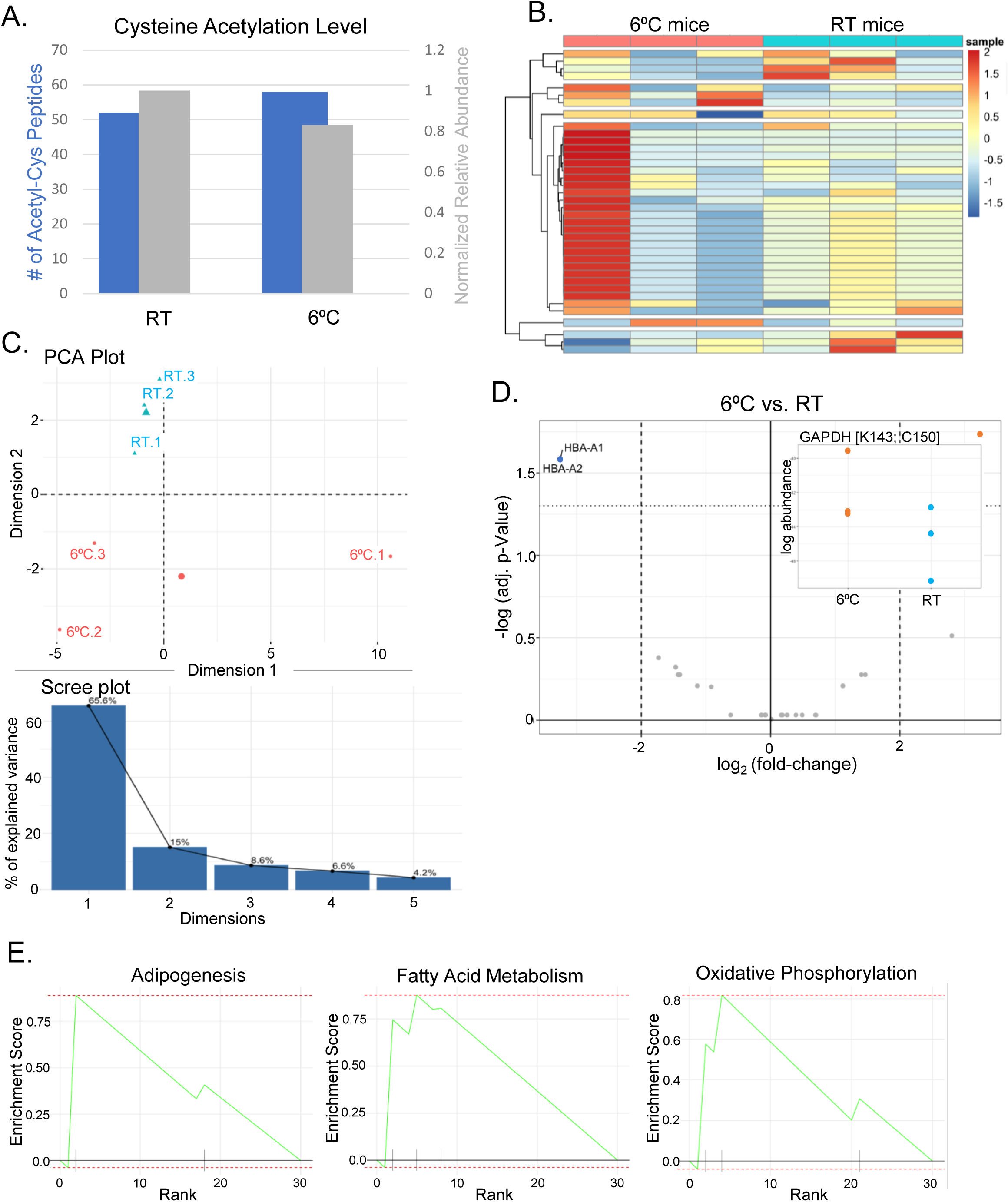
Cysteine acetylome in BAT from mice acclimated to 6□C compared to mice housed at RT. A. Overall abundance of the cysteine acetylome in BAT at RT and 6□C. In gray is the summed intensity of cysteine acetylated peptides, normalized to total peptide signal, and in blue is the number of unique S-acetylated peptides identified. B. Heatmap showing differences in the distribution of the cysteine acetylome in BAT from mice that were housed at RT vs mice housed at 6□C C. PCA of the cysteine acetylome in BAT from RT and 6□C mice. Scree plot showing the relative contribution of each of the first 10 components. D. Volcano plot showing fold-change in abundance of cysteine acetylation at sites acetylated in both the RT and 6□C condition. Cutoffs at 2x fold change and p<0.05. E. GSEA plots showing enrichment for Adipogenesis, Fatty Acid Metabolism, and Oxidative Phosphorylation in the cysteine acetylome of 6□C acclimated mice compared to RT housed mice.

Next, gene-set enrichment analysis (GSEA) was used to identify differences in the cysteine acetylome of 6°C-challenged mice compared to the cysteine acetylome of RT-housed mice. We found a significant enrichment of the adipogenesis pathway in the cysteine acetylome of 6°C housed mice (Figure 4E). The next two most enriched pathways were Fatty Acid Metabolism and Oxidative Phosphorylation. While this does not resolve the details of how cysteine acetylation may be involved in the cold response in BAT, these pathway enrichments suggest a possible involvement of cysteine acetylation in BAT metabolic reprogramming upon cold exposure.

### Cysteine Acetylation on GAPDH reduces its activity

Of particular interest, two proteins showed statistically significant changes in cysteine acetylation levels in cold-challenged mice compared to RT controls (Figure 4D). Hemoglobin A (HBA) showed decreased levels of cysteine acetylation upon 6□C exposure, while GAPDH showed increased acetylation in the active site upon cold exposure.

Interestingly, initial characterization of the liver cysteine acetylome also identified GAPDH as a top hit. Thus, further experiments were undertaken to characterize the effect of cysteine acetylation on GAPDH. The murine liver tissue cysteine acetylome showed acetylation of GAPDH on Cys150; a known active site residue of GAPDH (Figure 5C). Cys150 is known to have multiple PTMs including ADP-ribosylation,^15,16^, S-(2-succinyl)cysteine (aka succination),^17^ and nitrosylation.^18^ Nitrosylation of Cys150 has been shown to inhibit the metabolic activity of GAPDH and direct its nuclear localization.^19^ In the nucleus, GAPDH mediates an apoptotic response; however, its translocation to the nucleus and its function there can also be dependent upon acetylation.^20^ ^21^

**Figure 5.**
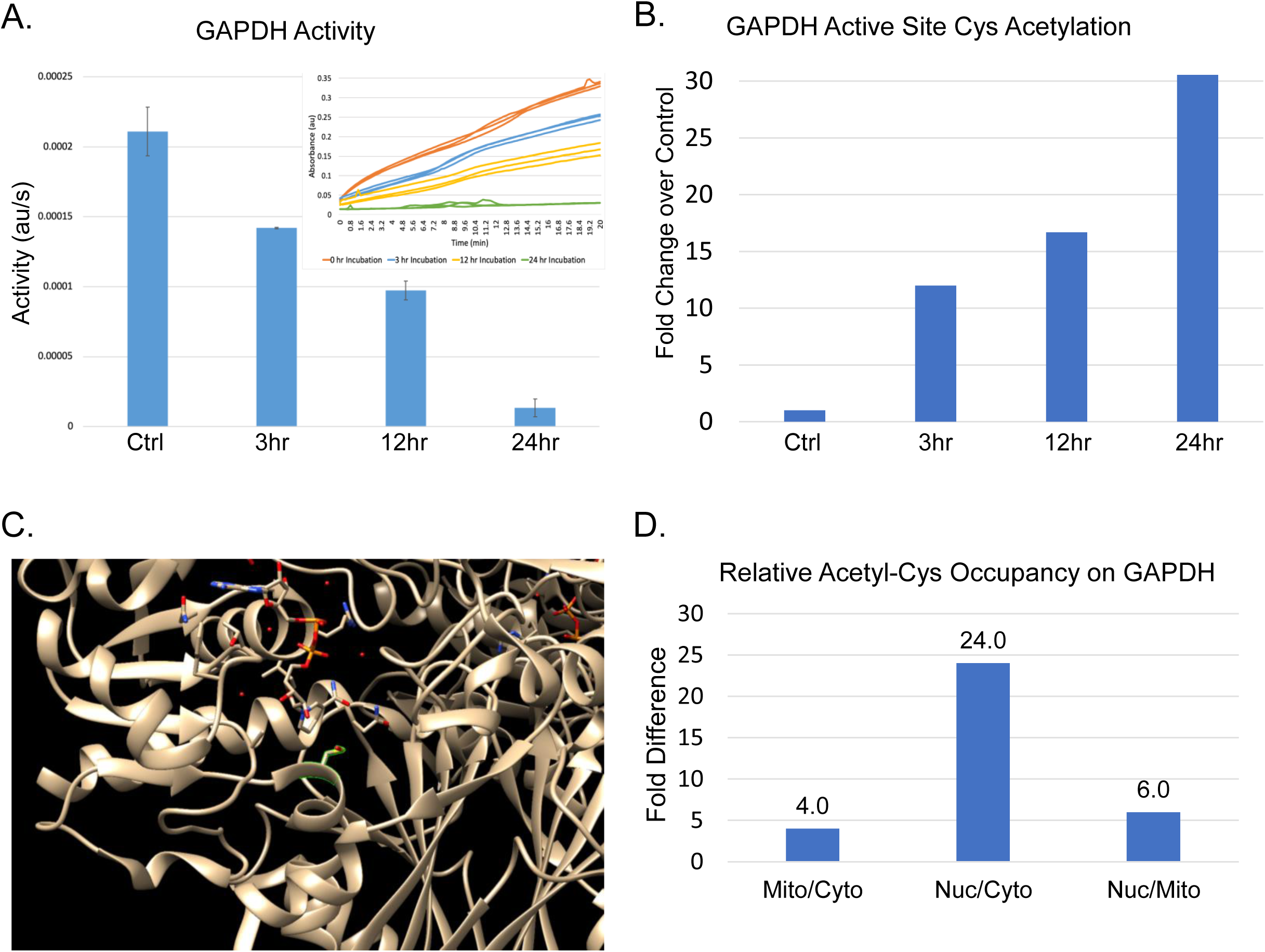
GAPDH acetylation and activity A. Activity of GAPDH as determined by *in vitro* assay upon increasing levels of cysteine acetylation by incubation with acetyl-CoA. B. Extent of GAPDH active site acetylation upon increasing length of incubation with 10mM acetyl-CoA. C. Active site structure of GAPDH with Cys150 highlighted. D. Relative fraction of GAPDH that is cysteine acetylated in the active site in different cellular compartments.

Thus, we prioritized further studies on the effects of cysteine acetylation on GAPDH activity *in vitro*, hypothesizing that modification of an active site cysteine residue might influence enzyme activity. To determine the effect of cysteine acetylation on GAPDH activity, we incubated purified GAPDH with acetyl-CoA. Enzymatic activity was quantified by an *in vitro* activity assay. GAPDH was incubated with 10 mM acetyl-CoA at 4□C for 0, 3, 12, and 24 hours. As a control, all samples were incubated at 4□C for 24 hours to ensure the only differences between the samples was the incubation time with acetyl-CoA. Following the 24-hour incubation period, each sample was divided into two. One portion was analyzed by MS to determine the relative level of cysteine acetylation on either of the two active site cysteine residues, and the other portion used to determine the activity of GAPDH by *in vitro* enzymatic activity assay. We found activity decreased with increasing acetylation of the active site cysteine residues (Figure 5A). After 24 hours of incubation with acetyl-CoA, active site cysteine acetylation had increased approximately 30-fold (Figure 5B), while the activity of GAPDH had decreased 90% compared to control (Figure 5A).

Because nitrosylation of Cys150 influences GAPDH subcellular localization, we inspected the subcellular distribution of cysteine-acetylated GAPDH. Interestingly, we found a unique cysteine acetylation signature of the S4R1W1 isoform of GAPDH. This isoform differs by a single amino acid substitution, A to S substitution at position 156 (A156S), from the reference isoform. Because of this single difference, only peptides that included residue 156 could positively be identified as belonging to one or the other isoform. Across all proteomic datasets collected, all active site peptides of S4R1W1 were acetylated on at least one residue, except only some samples that had been treated with DTT to reduce thioesters. In contrast, peptides with A156 were only sometimes acetylated. Furthermore, the subcellular fractionation studies showed localization of the S4R1W1 isoform to the nucleus. Despite overall levels of GAPDH being higher in the cytoplasm, the fraction of GAPDH acetylated was 28 times higher in the nucleus than in the cytoplasm and 18 times higher in the nucleus than in the mitochondria (Figure 5D). Analysis of the individual isoforms of GAPDH involved revealed that the enrichment of GAPDH active site cysteine acetylation in the nuclear fraction was driven by the localization of the consistently acetylated S4R1W1 isoform to the nucleus, not by an increase in acetylation of the canonical isoform. All S4R1W1 active site peptides were acetylated, while none of the canonical active site peptides observed in the nuclear fraction were acetylated. The lower levels of the canonical isoform in the nucleus increased the observed frequency of active site acetylation across both isoforms combined. Thus, the unique cysteine acetylation signature and nuclear localization of the S4R1W1 isoform of GAPDH suggest that this isoform may play a distinct role in nuclear processes, possibly related to the regulation of gene expression.

## DISCUSSION

Due to the presence of a reactive thiol, cysteine is an amino acid that plays a key role in enzyme catalysis. Beyond catalysis, cysteine side chains are frequently covalently modified, with succination^22^, nitrosylation^23^, and palmitoylation^24,25^ being some of the more common cysteine-based PTMs. Reactive oxygen species (ROS) are also mediated through post-translational modification of the cysteine side chain.^23,26^ In contrast to the rich landscape of lysine acylation, far less is known regarding acyl modifications to cysteine. Previous studies have only identified cysteine as susceptible to long-chain acylation, specifically cysteine palmitoylation, which is involved in protein transport and localization, and implicated in neurological and immune function.^25,27,28^

Cysteine also plays a key role in intramolecular transfer, notably in ubiquitin conjugation enzymes.^29^ Beyond this canonical role, the possibility of acyl PTM transfer from a cysteine side chain to lysine was first reported in 2017. *In vitro* studies showed that an acetyl moiety on a cysteine residue’s side chain can undergo intramolecular transfer to a lysine residue on the same peptide.^5^ This finding suggests a novel pathway for non-enzymatic lysine modification and raises the possibility of cysteine being modified by short-chain acyl groups, in addition to the well-documented long-chain acyl modifications.^25^

The reactivity of the thioester bond in Acetyl-CoA suggests that under physiological pH, a thioester exchange reaction could transfer a short-chain acyl group from acyl-CoA onto the cysteine side chain.^30^ A similar S-linked short-chain acyl modification has been reported in the active site of fatty acid synthase.^31^ Although this modification is not specifically on the cysteine side chain, it supports the possibility of S-linked short-chain acyl modifications. Despite these advances, there has been no successful identification or mapping of an S-linked short-chain cysteine acylome.

We here report the discovery of a novel side-chain PTM of cysteine – acetyl-cysteine. This modification expands the encyclopedia of acetylated residues and introduces a novel class of PTMs—short chain acylation of cysteine. The cysteine acetylome is distributed across the murine proteome and was confirmed by MS and by chemical ablation. This modification was previously unreported having evaded detection likely due to its instability under common sample preparation conditions. The thioester bond that attaches the acetyl moiety to the cysteine side chain is reduced by DTT^7^ and is sensitive to pH.^6^ This renders it susceptible to cleavage under standard MS sample preparation conditions. Furthermore, MS fragmentation approaches can also cleave the thioester bond of acyl-cysteine modifications.^32^

The absence of antibodies that successfully target acetyl-cysteine renders specific enrichment of the cysteine acetylome technically infeasible with currently available reagents. Similar modifications such as lysine acetylation have traditionally been enriched by antibody pull-down prior to characterization by MS. Identifying a labile, unenriched PTM presents difficulties that have heretofore prevented the identification and characterization of the cysteine acetylome.

Previous studies attempting to characterize/quantify long-chain acyl modifications to cysteine such as palmitoylation have often assumed that all acyl cysteine modifications consist of long-chain acylation. These studies have repeatedly identified many sites of modification that are not positively identified as being palmitoylated by MS. Short-chain acyl modifications to cysteine were not known in the literature and not searched for and hence could not be observed even if present.^24,33,34^ The presence of thioester bound acyl-cysteine sites that did not map to known sites of palmitoylation hinted at the existence of short-chain cysteine acylomes.

Lysine acetylation is primarily an enzymatically mediated modification; however, lysine acetylation can also occur non-enzymatically.^35^ At physiological pH, the thiol group of the cysteine side-chain is notably more reactive than the amine group of the lysine side-chain.^36^ Non-enzymatic acetylation of cystine would be expected to be more rapid and favorable than lysine acetylation. The presence of non-enzymatic modification of cysteine does not indicate the absence of enzymatically targeted modification. In this paper we detail *in vitro* results which indicate the presence of non-enzymatic modification of cysteine, especially at elevated concentrations of acetyl-CoA. This should not be construed as an indication that enzymatic modification of cysteine is not physiologically relevant. Indeed, differences in the reactivity of specific cysteine residues towards acetyl-CoA in lysate compared to purified protein samples could plausibly be interpreted to indicate the presence of enzymatic regulation of cysteine acetylation in lysate samples. Enzymatic acetylation of cysteine is a natural and obvious path to targeted cysteine acetylation, however the local microenvironment of a cysteine side chain can also function to target non-enzymatic acetylation of cysteine. The pKa of the cysteine side-chain can vary by two or more pH units depending on local microenvironment.^36^ This can cause order of magnitude differences in the reactivity of a cysteine side-chain towards acetyl-CoA at physiological pH. Binding of acetyl-CoA in proximity to a cysteine side-chain could also cause non-enzymatic targeting of specific cysteine residues.

Initial efforts to characterize the cysteine acetylome showed a widespread low-abundance acetyl-cysteine signature. The specific sites and proteins modified were inconsistent between samples. This could partially be attributed to the low abundance of the modification and the incomplete sampling of the MS. As a labile modification, cysteine acetylation would be expected to be susceptible to intramolecular transfer between modified and unmodified cysteine residues. Immediate alkylation of exposed cysteine side-chain residues upon thawing and lysis of tissues samples helped improve the reproducibility of the sites identified in the cysteine acetylome. Future work characterizing the cysteine acetylome will need to be cognizant of this scrambling of the cysteine acetylome during periods of sample preparation and manipulation.

Subcellular analysis of the cysteine acetylome revealed a preference for the cytoplasm over the nuclear and mitochondrial compartments. This was initially unexpected due to pathway analysis of the cysteine acetylome that showed enrichment of metabolic pathways, and the obvious role of the mitochondria in metabolism. This again points to the possibility that the role of cystine acetylation is not simply a function of metabolism. Rather, it seems that there is a physiological role involved in metabolic pathways, but not inherent to the processes of metabolism itself. Possible roles that fit this description would include protection of reactive cysteine side-chains or perhaps regulation of multi-functional proteins. The potential of cysteine acetylation to occur non-enzymatically also raises the possibility of it’s functioning as a non-enzymatic sensor of acetyl-CoA levels. In such cases one can envision a feedback system where cysteine acetylation regulates protein activity based on acetyl-CoA level.

Characterization of the cysteine acetylome in BAT from mice house at RT and mice cold adapted to 6□C did not show massive restructuring of the cysteine acetylome. Rather, there was one hit that showed a statistically significant increase in cysteine acetylation – GAPDH. One limitation of this method of comparison is that it required cysteine acetylation be identified in both conditions before comparison was possible. Hence sites that only showed acetylation in one condition were not included.

GAPDH had previously been identified as one of the top hits in the liver cysteine acetylome. GAPDH was acetylated on the active site cysteine Cys150. This was particularly interesting because nitrosylation of that residue was known to drive trafficking of GAPDH to the nucleus and trigger one of the many moonlighting functions of GAPDH. Nitrosylation of GAPDH was driven by exogenous addition of NO. It could be that in a physiological setting acetylation of Cys150 rather than nitrosylation is was drives nuclear translocation and the moonlighting functions of GAPDH. Consistent with this, the fraction of GAPDH that is cysteine acetylated in the active site is much higher in the nucleus than in the cytoplasm or mitochondria. While we have not directly observed trafficking of GAPDH to the nucleus as a result of acetylation, these results are consistent with such trafficking having taken place.

Acetylation of the GAPDH active site cysteine residues led to ablation of GAPDH activity. This was to be expected given that the cysteine residues are involved in catalysis. This raises the possibility that the acetylation of the active site cysteine in GAPDH may serve as a regulatory signal that not only inhibits the metabolic activity of GAPDH but may also be involved in regulating its localization. Given the multiple potential moonlighting roles of GAPDH in the cell this presents exciting new avenues for research into GAPDH activity and regulation

Given that the moonlighting roles of GAPDH do not involve catalysis of glyceraldehyde-3-phosphate, cysteine acetylation may serve to control which role GAPDH takes on in the cell. Multi-tissue survey of the cysteine acetylome showed acetylation of the multiple metabolically relevant proteins across different tissues. Cysteine acetylation may inhibit the activity of those proteins like it does GAPDH. Cysteine acetylation can block the reactive side chain of cysteine thereby preventing both disulfide bonding and catalysis in proteins dependent on cysteine for either their structure or function. However, there are several other possibilities as well. Cysteine acetylation could serve as a protecting group preventing undesired side-reaction with the cysteine side chain during translation or translocation of proteins. This may be the case for GAPDH inhibition of its metabolic active site seems to be key to both its translocation and its nuclear activity, however this inhibition may serve not just a signaling role, but also to directly prevent unwanted catalysis from GAPDH when serving in a moonlighting role.

A major limitation to further exploration of the cysteine acetylome is the absence of effective tools for acetyl-cysteine enrichment and visualization. The effectiveness of future characterization of the cysteine acetylome will be in large part dependent on the development of an effective antibody or other means of enrichment. Previous studies characterizing the long-chain cysteine acylome have used a variety of chemical approaches to achieve enrichment.^24,33,34^ Unfortunately, these approaches all fail to differentiate between long-chain and short-chain acylation. The ability to either enrich for specific S-acyl modifications, or match specific enriched sites to specific acyl modifications will be key to better understanding of the physiological role of short-chain acyl modifications of cysteine.

In conclusion, we describe the identification of acetyl-cysteine as a widespread protein post-translational modification. The presence of cysteine acetylation immediately raises the possibility of other acyl-CoA derived short-chain acyl modifications to cysteine. Lysine is known to have 27 different acyl modifications derived from acyl-CoA species, of which 21 are short-chain acyl modifications.^1,37^ It is not unreasonable to posit that cysteine may have a similar suite of short-chain acyl PTMs beyond the cysteine acetylome described here awaiting discovery.^23^

## Supporting information

Table 1. TCEPvDTT

Table 2. EarlyAlkylation

Table 3. LiverTopSites

Table 4. Liver+AcetylCoA

Table 5. TissueSurveyTopSites

---

**Figure S1.**
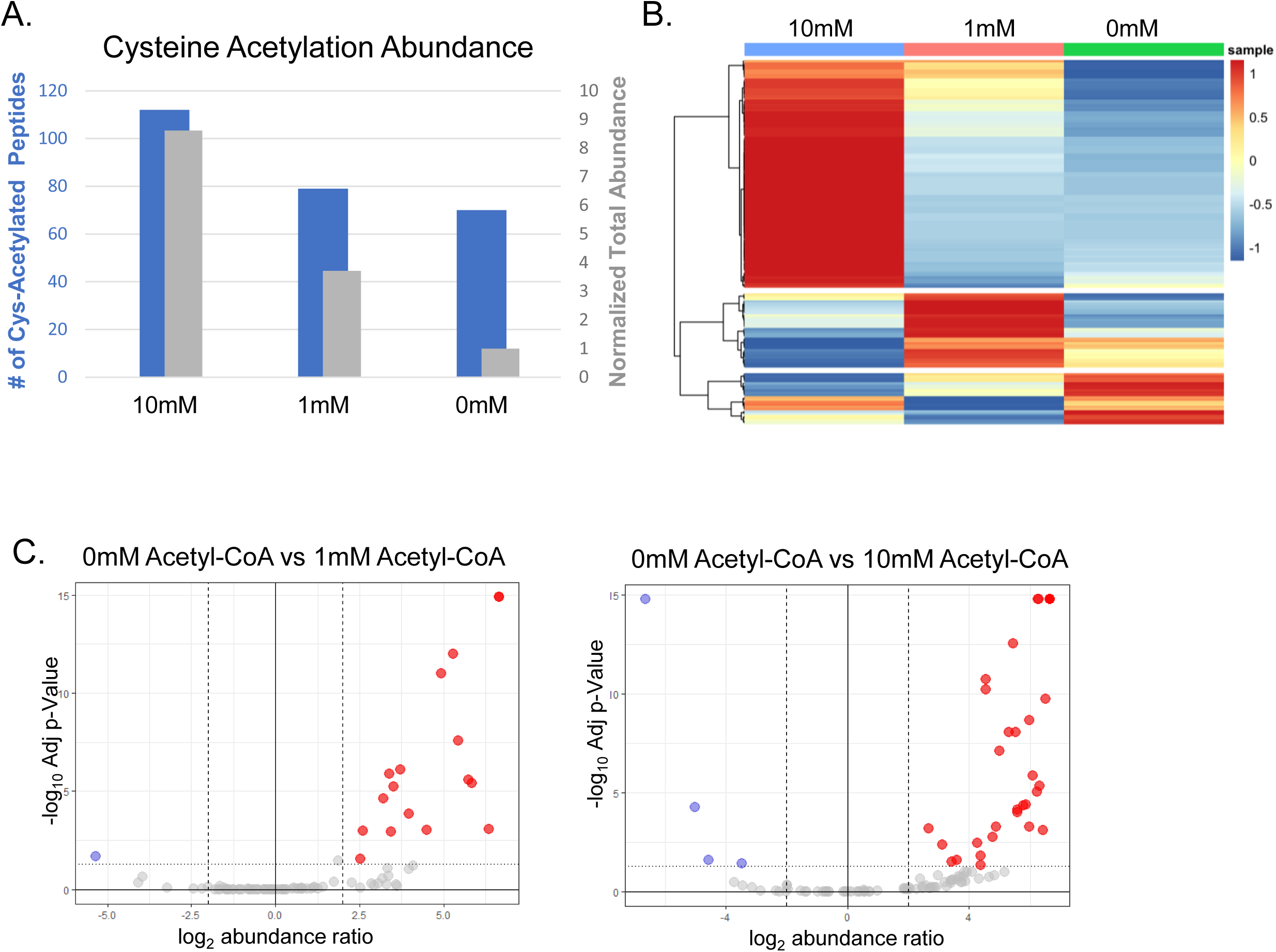
Incubation with Acetyl-CoA increases cysteine acetylation. A. Heatmap showing distribution of cysteine acetylation in samples incubated with acetyl-CoA at 10mM, 1mM or 0mM (control). B. The summed intensity of cysteine acetylated peptides, normalized to total peptide signal, (in gray) and the number of unique S-acetylated peptides identified (in blue) for each treatment condition. C. Volcano plots showing the relative abundance of cysteine acetylation in samples incubated with 1mM or 10mM acetyl-CoA compared with control sample incubated with 0mM acetyl-CoA.

**Figure S2.**
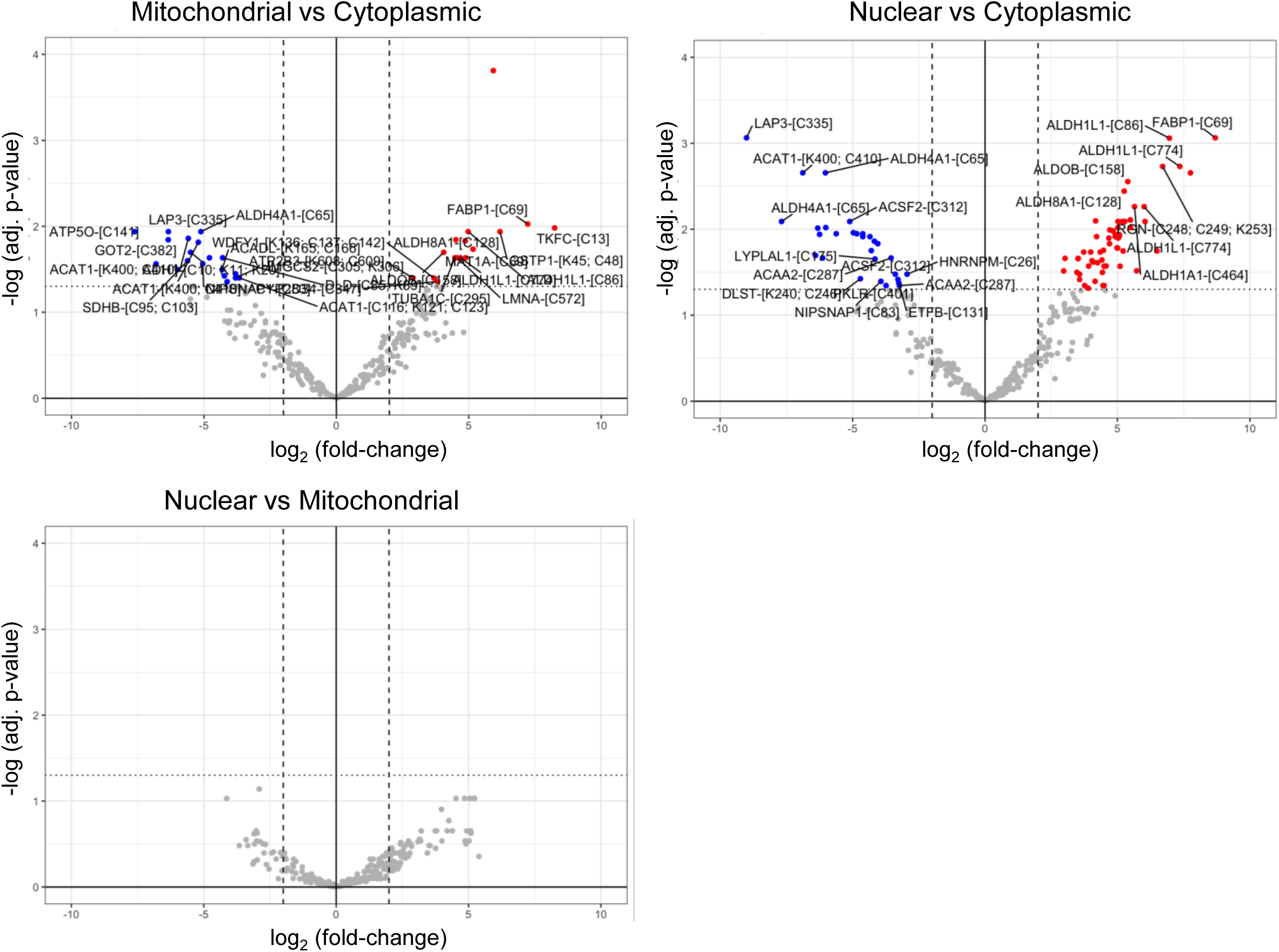
Volcano plots showing proteins and sites with the greatest differences in modification level between subcellular compartments.

## METHODS

### Animals

Ethical approval was obtained from the Institutional Animal Care and Use Committee (IACUC) at Duke University. The approved protocol is registered as Protocol Registry Number A035-23-02, titled “Modifications of Mouse Genes for Metabolic Research”. All animal experiments and procedures were conducted in strict accordance with the approved IACUC protocol and adhered to the guidelines and regulations set forth by the Animal Welfare Act and the Guide for the Care and Use of Laboratory Animals. All mice used in this study were wild type C57/Bl6J. Mice were housed in 12:12 light dark cycle facility and fed on standard chow.

### Method Details

Tissue Collection and Preparation. Mice were euthanized, and tissues were immediately removed and flash frozen by clamping with pliers pre-chilled in liquid nitrogen. Frozen tissues were stored at -80□C until further processing. Tissues were pulverized using a mortar and pestle chilled in liquid nitrogen and stored at -80□C until use. Pulverized tissue was weighed out into an Eppendorf tube chilled in liquid nitrogen to prevent thawing during the weighing process.

Tissue Lysis and Protein Extraction. Tissue powder was thawed by adding Lysis Buffer (5% SDS, 50mM Tris pH 7.4, 25mM TCEP, 20mM Iodoacetamide (IAC)) at a volume of 130μL per 20mg of tissue powder. IAC was freshly prepared from powder immediately prior to each experiment, and TCEP and IAC were added to the lysis buffer immediately before use. Samples were thoroughly vortexed to ensure complete lysis and then incubated in the dark at room temperature for 20 minutes to allow for the alkylation of exposed cysteine residues. Following incubation, samples were centrifuged at 20,000g for 15-20 minutes at 16□C to pellet the genomic DNA. In cases where the DNA did not pellet effectively, samples were sonicated to shear the DNA before repeating the centrifugation step. The supernatant containing the proteins of interest was collected, and the pellet was discarded.

Sample Preparation using S-Trap Micro Columns. For each sample, 25μL of the supernatant was processed using S-Trap micro columns. 12% phosphoric acid was added at a ratio of 1:10 to the 25μL sample (2.5μL of 12% phosphoric acid per sample), followed by the addition of 165μL of S-Trap buffer (90% methanol with 100mM TEAB, pH ∼7). The entire sample was loaded onto an S-Trap micro column placed in an Eppendorf tube and centrifuged at <4000g for <3 minutes. Columns were washed 4 times with 150μL of S-Trap buffer (90% methanol, 100mM TEAB, pH ∼7) and centrifuged at <4,000g for <3 minutes at 16□C, discarding the flow-through after each spin. An additional spin was performed after the final wash to ensure complete removal of residual S-Trap buffer.

Trypsin Digestion. Each S-Trap micro column was transferred to a fresh Eppendorf tube, and 20μL of Trypsin solution containing a 1:10 (w/w) ratio of protease to protein was added directly onto each micro column. Pre-prepared Trypsin samples containing 20μg in 40μL total volume were used, with 20μL (10μg) of trypsin solution added to each column. Samples were incubated at 37□C for 2.5 hours.

Peptide Elution and Preparation for Mass Spectrometry. Peptides were eluted from the S-Trap micro columns in three steps: first with 40μL of 50mM TEAB pH 8 (added directly to the trypsin in the S-Trap), then with 40μL of 0.2% formic acid, and finally with 40μL of 50:50 ACN:H2O with 0.1% formic acid. The eluate was frozen in liquid nitrogen and dried overnight by speed vacuum. Dried samples were resuspended in 100-200μL of 0.1% formic acid, vortexed to aid in resuspension, and incubated at room temperature to allow for complete resuspension. Samples were then centrifuged at 20,000g for 10 minutes at 4□C, and the supernatant was collected.

Peptide Quantification and Mass Spectrometry Analysis. The peptide concentration of the supernatant was quantified using the Pierce Colorimetric Peptide Assay (ThermoFisher 23275). The supernatant was then transferred to mass spectrometry vials for loading and characterization by Q-Exactive Mass Spectrometry. All samples were stored at -80□C until analysis.

### Subcellular Fractionation

Mouse liver tissue was obtained as described previously. Tissue was fractionated by differential centrifugation at 70g, 700g, and 7,000g into nuclear, cytoplasmic and mitochondrial fractions.

### 6LJC vs RT BAT

Age-matched mice were housed at either 6□C or at RT. Mice were cold-adjusted by housing at 6□C for four weeks. BAT was harvested and processed as previously described for liver tissue.

### GAPDH in vitro acetylation

Purified GAPDH from rabbit erythrocytes commercially available from Sigma (G2267). GAPDH at 1mg/mL was acetylated by incubation with 10mM Acetyl-CoA in PBS at 4□C.

### GAPDH activity assay

GAPDH activity was measured using the commercially available GAPDH Activity Assay Kit, (Sigma, MAK277).

### Statistics

Hierarchical clustering and heatmap plotting were performed using the pheatmap R package (version 1.0.12) with correlation as the distance metric and the “average” as the clustering method. Principal Component Analysis (PCA) was performed on scaled and centered data using the R packages FactoMiner (version 2.11)^38^ and factoextra (version 1.0.7).^39^ Differential abundance analysis was performed using the limma R package (version 3.58.1).^40^ Gene set enrichment analysis (GSEA) was performed using the fgsea R package (version 1.29.1).^41^ Data wrangling and visualization were performed using the tidyverse collection of R packages (version 2.0.0). Analyses were perfomed in R (version 4.3.2).

## ACKNOWLEDGEMENTS

We would like to thank Dr. Rana Gupta and Lavanya Vishvanath for providing mouse BAT samples, and Dr. Derek Zachman for reviewing an early draft of the manuscript. We would like to acknowledge funding support from the National Institutes of Health and the NIA grant R01AG045351, R01DK115568, and R21AG080334 (MDH). E.K.K. was supported by grant 3R01DK11556803S1, and is supported by an NIH/NIGMS training grant to Duke University Pharmacological Sciences Training Program (5T32GM007105-40). The content is solely the responsibility of the authors and does not necessarily represent the official views of the National Institutes of Health or other funding sources.

## DATA AVAILABILITY

Proteomic data is deposited on Proteome Exchange under PXD052367.^37^

## CODE AVAILABILITY

Code used to analyze data available at https://github.com/hirscheylab/Cysteine-S-acetylation-is-a-post-translational-modification-involved-in-metabolic-regulation/tree/main.

Datasets used for analysis included as Supplemental Datasets 1-48.

## AUTHOR CONTRIBUTIONS

E.K.K. Conceptualization, Methodology, Investigation, Data curation, Writing - original draft. A.B. Software, Formal analysis, Visualization. Y.L. Investigation. P.A.G. Resources, Investigation, Writing - Review & Editing. M.D.H. Supervision, Funding acquisition, Writing - Review & Editing.

